# Evidence for Early European Neolithic Dog Dispersal: New Data on South-Eastern European subfossil dogs from Prehistory and Antiquity Ages

**DOI:** 10.1101/609974

**Authors:** Miroslav Marinov, Iskra Yankova, Boyko Neov, Maria Petrova, Nikolai Spassov, Peter Hristov, Georgi Radoslavov

## Abstract

**Objectives:** The history of dog domestication is still under debate, but doubtlessly, it is a process of an ancient partnership between dogs (*Canis familiaris*) and humans. Although data on ancient DNA dog diversity are scarce, it is clear that several regional dog populations had been formed in Eurasia up to the Holocene. During the Neolithic Revolution and the transition from hunter-gatherer to farmer societies, followed by civilization changes in the Antiquity period, the dog population structure also changed. This process was due to replacement with newly formed dog breeds.

**Methods:** In this study we have presented for the first time mitochondrial data about South-Eastern Europe (the Balkans) ancient dog remains from the Early Neolithic (8 000 years BP) to the Late Antiquity ages (up to 3th century AD). A total of 25 samples were analyzed using the mitochondrial D-loop region (HVR1).

**Results:** The results have shown the presence of A (70%) and B (25%) clades throughout the whole investigated period. In order to clarify the position of our results within the ancient dog population in Eneolithic Eurasia, we performed phylogenetic analysis with the available genetic data sets. This data revealed a similarity of the Bulgarian dogs’ structure to that of ancient Italian dogs (A, B, and C clades), which suggests a new prehistoric and historic Mediterranean dog population. A clear border can be seen between South-European genetic dog structure, on the one hand, and, on the other hand, Central-West (clade C), East (clade D) and North Europe (clades A and C). This corresponds to genetic data for European humans during the same period without admixture between dog populations.

**Conclusions:** Our data have shown for the first time the presence of clade B in ancient Eurasia. This is not unexpected as the B haplogroup is widely distributed in extant Balkan dogs and wolves. The presence of this clade both in dogs and in wolves on the Balkans may be explained with hybridization events before the Neolithic period. The spreading of this clade across Europe together with the A clade is related to the possible dissemination of newly formed dog breeds from Ancient Greece, Thrace and the Roman Empire.

## Introduction

In recent years, it has been increasingly assumed that the dog’s domestication was a very early process that began at the end of the Pleistocene. The dog’s domestication is apparently linked to the gradual synanthropy of wolf populations as a result of commensal relations between man and wolf at the end of the late Pleistocene [1,2]. The first data from dog-like remains came from the Razboinichya Cave in the Altai Mountains of Siberia (33 000 years BP) and the Goyet cave in Belgium (c. 31 700 BP) [3,4]. Genetic studies of these remains have not shown any similarity to recent wolves and dogs [3,5]. The Pleistocene domestication cannot be accepted as absolutely certain, given the large morphological variability of Pleistocene wolves [6].

Therefore, we can assert that signs of safe substitution are found only in the borderline between the Paleolithic and the Mesolithic. Since then, the main questions have been related to the place and time of origin, domestication and influence of hybridization events between domesticated dog and local wolf populations.

According to mitochondrial DNA (mtDNA) studies, there are six main clades, assigned A, B, C, D, E, and F, and many sub-clades, characterizing dog population in the world [7–9]. Clades A, B, and C are the most widely distributed (95.9%) among recent dog populations, while D, E, and F have regional geographic distribution [10]. For example, in West Eurasia, the sub-haplogroup A1 is with a frequency of about 70%, while clades B and C are present with frequencies of 20% and 10%, respectively [7,9]. In the Middle East, this proportion is almost the same but with more worldwide presented sub-haplogroups [11].

Analysis of ancient dog DNA is still scarce and does not include many geographic regions. Despite this, available data reveal quite different population structure concerning present day dogs. These alterations are due to the emergence of the first civilizations as well as the early historical and modern human migration. For example, pre-Columbian American dogs were identical to East-Eurasian (Siberian) dogs which are nowadays mixed with West-Eurasian dogs, after the Age of Discovery (15th century) [5,12,13]. Similarly, in the Pacific, including the islands of Polynesia, a post-Lapita dog introduction from southern Island Southeast Asia has been suggested [14].

### Ancient dog populations from Europe

One of the most investigated regions concerning ancient dog mtDNA is Europe. Up to date, most studies have shown that old Europe (from the Pleistocene to the Holocene) may be divided into at least three or four regions of dog populations based on mtDNA analysis. One of them is Central and West Europe, including France, Germany, Switzerland, and Hungary. Typical for these parts of Europe is the prevalence of clade C (over 80%) mixed with clade D [5,13,15,16] (Fig 1a). The second region includes Eastern Europe (Romania, Moldova, Ukraine) up to Iran and the Middle East. The basic clade in these regions is D (over 90%), and there are traces of A and C [5,13,16,17]. In Northern Europe (Scandinavia and Estonia) there are clades A and C [5,16,18]. Also, the most presentable nowadays clade D is missing [7,19,20]. There has been only one study from the Mediterranean region concerning ancient dog populations [21]. This research includes only five samples (three wolves and two dogs), and the obtained results have shown the presence of clades A, B, and C [5,21]. Most of the authors have suggested that clade A is associated with Neolithic farmer migration in ancient Europe from the Middle East, while clade B probably originated from Southeastern Europe (the Balkans and the Apennines) because of the high frequency of extant Balkan wolves with similar haplotypes [5,21–24].

**Figure 1.**
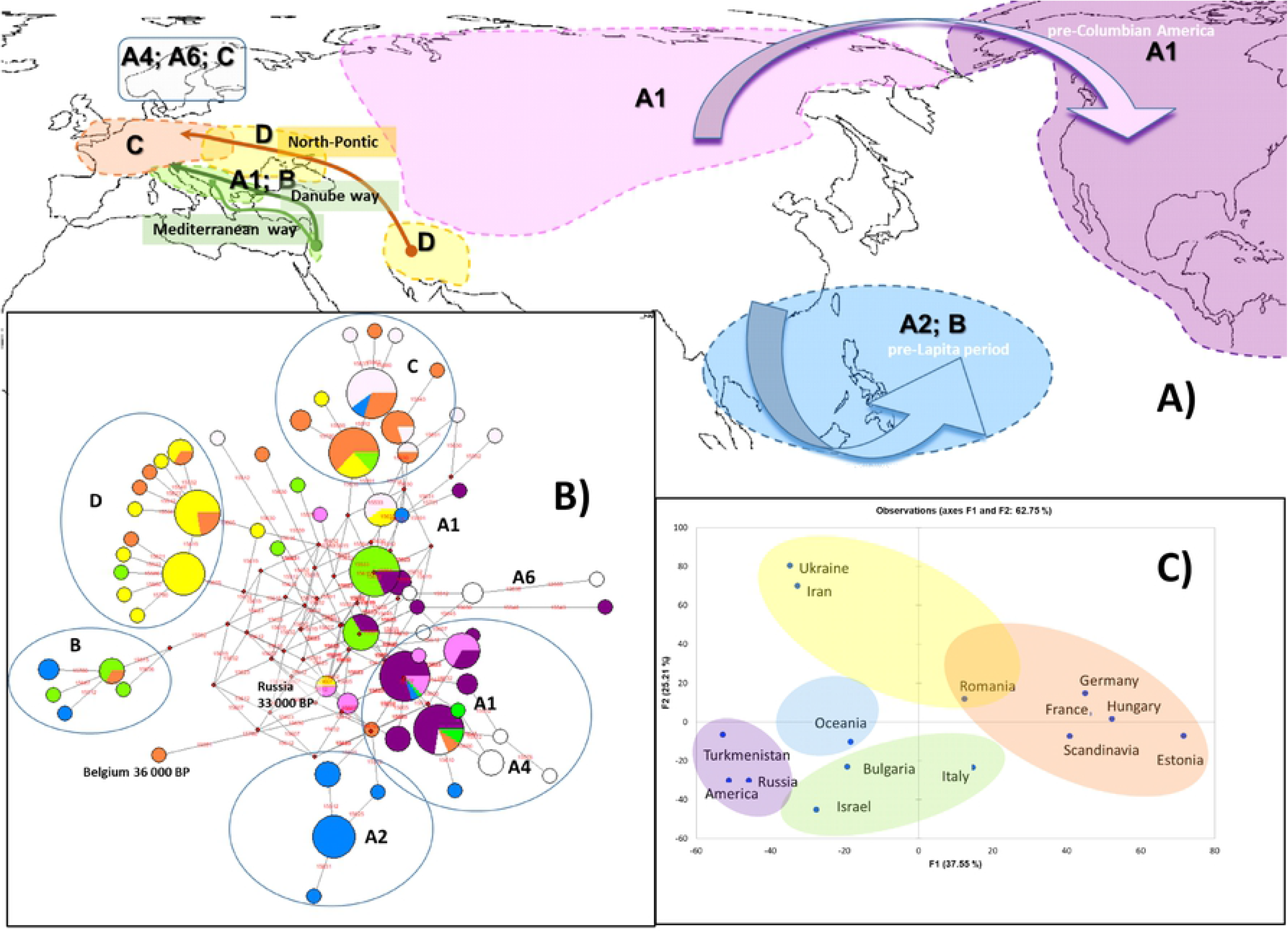
Population structure of ancient dogs worldwide based on mtDNA data. a) A world map showing distribution of main haplogroups in ancient dog. The main migration routes are also shown. b) The reduced median network of the main mtDNA haplotypes from ancient dogs of different regions of the world, c) Principal components analysis of the main mtDNA haplotypes from ancient dogs of different regions of the world

All these data from pre-historic Europe have shown a dramatic difference between ancient and present day dog populations. This change may be explained with the replacement of local dog breeds with more improved breeds from the South part of Europe, the Mediterranean region, and the Near East during the Late Antiquity and the Medieval ages [21].

### Ancient dog populations in Central Asia and pre-Columbian Americas

Another well studied area of research is related to the region of pre-Columbian America and the migration of Paleolithic hunter-gatherers and their dogs from Siberia via the North Asian Mammoth steppe via the Beringia land bridge roughly 14 000 years ago [25,26]. MtDNA studies of ancient dogs from Siberia and the Americas reveal a similar genetic profile, i.e. the presence of specific A1 and A2 subclades [5,12,16,27,28] (Fig 1a). A similar mtDNA profile with a prevalence of clade A has been found in ancient dogs from Turkmenistan (Central Asia) [16].

### Ancient dog populations in South-Eastern Asia and the Pacific

The third most studied geographic regions are South-Eastern Asia and the Pacific Islands, including Australia (Fig 1a). Characteristic of these regions is the Neolithic pottery Lapita culture, where a Pacific Ocean people flourished in the Pacific Islands from about 3 600 BP to about 2 500 BP from continental Asia. A basic characteristic of dog populations in this region is the prevalence of sub-clade A2 and a few cases of B and A4’5 [13,14,29]. These sub-haplogroups are typical for recent South-Eastern Asia but with different frequencies [7,8]. Interestingly, the sub-clade B2 is different from West Eurasian dog’s B1 mytotypes [9].

In conclusion, dogs are a very interesting object, because of the direct evidence of historical human migration processes. However, there are still many uninvestigated geographic regions. White spots of ancient dog genetic diversity are, for example, the Mediterranean region, including South Europe, the Near East, and the Fertile Crescent, and also other domesticated centers in Central and East Asia (China and India), etc.

Our study tries to enrich data for ancient South-Eastern Europe – the Balkan Peninsula (Bulgarian) dogs, based on mtDNA (partial D-loop region, HVR1) analysis. We have performed phylogenetic analysis to compare our results with other ancient and recent European dog and wolf populations.

## Results

### Phylogenetic analysis and haplogroup classification

Ancient DNA was successfully amplified in 21 out of 25 samples tested. Out of the 21 sequences, 16 were used for phylogenetic analysis, because only they covered informative sites for haplogroups assignment. From them, eleven sequences were assigned to clade A (68.7%), four belonged to clade B (25.0%), and one to clade D (6.2%). These sequences have informative sites from position 15 595 bp to 15784 bp (189 bp) of the first amplified region of HVR1 according to ref. sequence EU789787 [8]. We analyzed only this region, because most of the ancient dog sequences available in GenBank also used this region as the most informative one for haplogroups assignment. Four of our not assigned sequences have only the third amplified fragment of HVR1, which is not declarative concerning haplogroup assignment (S1 Table).

All analyzed sequences in this study belonging to clade A and B are equally dispersed in investigated regions in Bulgaria as well as in historical time. In contrast, the highly dispersed in ancient East Europe clade D was found only in one sample from the Late Antiquity period (S1 Table).

### Comparative analyses of ancient dog haplotypes and Bulgarian samples

About 230 ancient dog sequences available in GenBank and Dryad Digital Repository package [17,40] were explored by Reduce-median network analysis. In this analysis, ancient dog samples were from ancient Eurasia, pre-Columbian America and Pacific pre-Lapita period (Fig 1b; S1 Table). All haplotypes were separated into four main clades: A, B, C and D. There is a clear geographical differentiation between the distribution of C and D clades, which are specific for Central, West and East Europe. Clade A is split into many sub-clades, i.e. A2 for South-Eastern Asia and the Pacific, A1 for Siberia and Central Asia, and pre-Columbian America, A4 and A6 for North Europe (Scandinavia), and A and B clade are characteristic for the Mediterranean region (Italy, Bulgaria and Israel). Similarly, PCA analysis grouped all ancient samples into five distinct clades (Fig 1c).

### Comparative analysis among ancient and recent Balkan dogs as well as recent gray wolf haplotypes distribution

About 120 haplotypes including recent Balkan dogs and wolves populations as well as ancient Bulgarian and Italian samples were analyzed by Median-joining network (S1 Fig). All samples (dogs and wolves) were explored as described previously from position 15 595 bp to 15784 bp (189 bp) of the first amplified region of HVR1 according to ref. sequence EU789787 [8], and haplotypes were determined according to the classifications of [9] and [34] as well as MitoToolPy program [35]. The results show that most of the ancient and recent dog haplotypes from clades A and B have similar or identical to recent Balkan wolves haplotypes. The Balkan wolves haplotypes are not grouped into C and D clades. Moreover, there are wolves haplotypes which not include present-day dog haplotypes.

## Discussion

### Balkan Neolithic and Chalcolithic periods: specifics of human societies and understanding of early European farmer migrations ways from the Near East

The South-Eastern Europe includes the Balkan Peninsula and the Apennines. These geographical regions are important points concerning human history and migration processes as well as the accompanying domesticated animals and the crops from the Near East. It is well known that during the Last Glacial Maximum (22 000–14 000 BP) the sea level of the World Ocean was about 125 meters lower than nowadays [41]. Therefore, large territories of land were connected. For example, the Balkan Peninsula was connected terrestrially to Anatolia via the Sea of Marmara in the east and also to a large part of the Apennines via the Adriatic Sea in the west [21]. This enormous contact zone was a precondition for the free movement of human societies but also of wild animal populations during the Mesolithic the and Early Neolithic periods.

Therefore, Early Neolithic cultures from the Fertile Crescent and Anatolia were directly connected to South-Eastern Europe (especially during the Late Neolithic period 8 000 BP). During this period three migration ways for dissemination of Neolithic farmers into Europe existed – the Mediterranean, the Balkans (the Danube River) and the North Pontic steppes (Fig 1a).

The most recent study on human population from the Mesolithic to the Iron Age has revealed that human populations on the Balkans (South-Eastern Europe) and in Anatolia shared a similar genetic profile in contrast with North Pontic Steppe Neolithic humans and those in Central and West Europe [42]. Despite the similarity of the Eastern and the South-Eastern Neolithic and Chalcolithic cultures, these genetic data suggest different migration routes (the Mediterranean, the Danubean and the North Pontic) of farmers from the Fertile Crescent. The data of this research clearly show human migrations from North to East direction on the Balkans during the Chalcolithic and the Bronze Ages, but also dissemination of South-Eastern human farmers into the Central Europe region (LBK culture – Austria, 5100–5000 BC). This evidence also suggests different migration routes for livestock like cattle, goats, and sheep [35,43].

In the Neolithic to the Chalcolithic Age livestock farming on the Balkans was related related to intensive cattle, goats and sheep breeding [31,44,45], with a prevalence of cattle (over 30%) in the North Balkans and domination of sheep breeding in the south. This animal husbandry was accompanied everywhere by the presence in the studied settlements of ancient races of dogs, which in later epochs passed into specially selected breeds.

The processes of domestication and dog breed creation continued later up to the Early and the Late Antique periods. At this time the first data on dog breeds on the Balkans were described by Aristoteles [46] and later, in the Roman empire, by Xenophon (Cynegeneticus, 2,000 years ago). The authors commented on dozens of different dog breeds, including guard, hunting, and companion dogs.

### Population structure of ancient dogs from the Neolithic period

The question of dog origin is geographically, genetically, and archaeologically complex. Ancient DNA analysis from relevant areas of the world should allow better understanding of the evolution of dogs from their predecessor, the gray wolf. In this respect, there are a few hypotheses about dog domestication and spreading. Based on recent mtDNA haplotypes distribution, the common opinion is that in the center of domestication the genetic diversity is the highest, thus there must be a high level of various haplotypes from different clades. Due to this reason, the East Asian and the Near East (Anatolian and Fertile Crescent) centers were proposed for dog origin centers [8,11]. The migration routes from the centers of domestication are characterized by a bottleneck type of dissemination which is unique for bordered regions worldwide like Europe, North Africa, Pacific Islands etc. For example, the genetic profile of recent European dogs is characterized by the prevalence with various A1 and B1 sub-clades [9,47], while dogs in South-West Asia (Anatolia and Fertile Crescent) have a mixed profile with other A and B sub-clades [11]. Recent European C1, D1 and D2 subclades are considered as regional specific sub-clades and as a “relict” from ancient time for dog-wolf hybridization [7,8].

In contrast to molecular data about recent dogs, evidence for ancient dogs, though still scarce, showed homogeneous and simple genetic structure of prehistory dog populations (Fig. 1a). In West, Central and North Europe there is a prevalence of C clade, in East Europe and the Middle East (Iran) – of clade D, in Southeastern Asia, Australia, and the Pacific Islands – a prevalence of the specific sub-clade A2, and in Siberia, Central Asia (Turkmenistan), and pre-Columbian America – of specific A1 sub-clades [5,12–16,27,28] (Fig 1b). Key geographic regions for understanding dog history like South Europe and the Near East, North Africa, and ancient Central and South Asia (China and India) are still uninvestigated.

The prehistoric Neolithic Balkan dogs were primitive in terms of stage of domestication, as their morphological features were relatively close to the ancestral wolf morphology as the “*vlasac*” type [30,31,48]. Our data have shown a prevalence of dog A and B clades from the Neolithic to the Antique period. During this time, the proportion of these clades was preserved with minor changes. Despite the small size of the investigated samples, these results are in direct contrast with the border regions in the north direction where the Danube River serves as a border region. The investigated ancient dog samples from Romania, Moldova, and Ukraine (9 000 – 5 000 BP) showed a prevalence of C and D clades [17], (S1 Table). Otherwise, our data correlate with ancient samples from Italy and Israel, where, despite the small size of samples, a prevalence of clades A and C for Italy has been revealed [5,16,17,21]. These data, even insufficient, have suggested that A clade dog populations replaced old Central European ones. It is possible that this genetic profile is associated with the Fertile Crescent and Anatolian ancient dogs.

### Clade B, a possibility of South-Eastern European (Balkans) origin of dogs from local wolves predecessors

In modern dogs, clade B is widely distributed with a frequency of over 20% [7,9]. This clade consists of two sub-clades, B1 and B2. The B1 sub-clade is disseminated worldwide with a frequency of about 21%, while the B2 sub-clade is with regional distribution mainly in Eastern Asia with a frequency of about 10 % [9,47]. Although there is a high frequency of clade B in modern dogs, in ancient samples up to date, these subhaplogroups have been observed very rarely, about 1 %. There is one sample in France and Turkmenistan, and there are a few samples from South-Eastern Asia and Oceania [14,16,17]. In our sample sets B1 sub-clade is with a high frequency (over 20%). This haplogroup is common among the Neolithic, the Chalcolithic and the Bronze Age samples but is also present in the Antique period with almost equal frequency.

These data are not very surprising because clade B is widely and unusually distributed in recent Balkan wolf population, identified in several researches [5,22–24,37]. These studies proposed that clade B possibly originated from the Balkans, although there are some cases of B type wolves worldwide. In a previous study of Bulgarian native dogs, clade B was observed with about 20 % [38].

The only investigation concerning South Europe about ancient dogs and wolves has been carried out by [21 In this study they investigated three wolves or large canids from 14 000 – 10 000 years ago and two medium size dogs from 4 000 years ago. One of the wolves’ samples was assigned as clade B (PIC-2), while others belonged to A (PIC-3) and C (PIC-1) clades (S1 Fig). The authors hypotesized about the possibile origin of B clades from the Balkans due to the high frequency of B type Balkan wolves. In addition, the authors defined that in South-Eastern Europe (the Balkans and the Apennines) there is a high frequency of clade B in gray wolf population. The discovery of clade B in the primitive dog race (intermedius-vlasac morpho-group) from the Early Neolithic of Slatina (Sofia) presented new arguments in favor of the morphologically [30,48] and genetically [21] argumented hypothesis that some prehistoric to recent dog races (races which have the B-clade) originated on the Balkans.

Taken together, all data about ancient dogs’ populations up to now, even though partly insufficient, have revealed a massive replacement of ancient local dogs before the Antique period and nowadays. These processes may easily be explained with the creation of new dog breeds in South Europe during the Antique and the Roman Empire periods, which disseminated and replaced local dogs in north direction in Medieval Europe. A similar process was also observed in different time and geographic regions during the Ages of Discovery, where C and B sub-clades changed the genetic structure of local American dogs via the introduction of extant European dog breeds.

## Conclusion

In conclusion, our data have enriched the information about ancient dogs’ structure, especially in South-Eastern Europe. The results from the Neolithic, the Chalcolithic and the Antique periods in Bulgaria, have demonstrated the dominance of A (70%) and B (25%) dog clades and homogenic structures of these haplogroups throughout the whole investigated period but an absence of clade C and only one case of D clade (late Antique period). These data have revealed the similarity of the Bulgarian dogs’ structure to that of ancient Italian dogs (A, B and C clades). Also, our data have shown for the first time the presence of clade B in ancient Eurasia. This is not unexpected because of the fact that the B haplogroup was widely distributed in extant Balkan wolves and dogs. The presence of this clade both in wolves and in dogs on the Balkans may be explained with hybridization events before the Neolithic period. Spreading of this clade across Europe together with the A clade is related to possible dispersal dissemination of newly formed dog breeds from Ancient Greece, Thrace and the Roman Empire.

## Materials and Methods

### Archeological samples collection, ancient DNA isolation and PCR amplification and sequencing

Twenty-five samples (bones and dental material) from ancient dogs from the collection of NMNH-BAS were studied from several archaeological sites: Early Neolithic 8500-7500 BP (Gradeshnitsa/Malo Pole, Ohoden–Valoga Slatina, Sofia district); Late Neolithic 7500-7000 BP (Topolnica, Promachon and Budzhaka-Sozopol); Early Chalcolithic 6950-6500 BP (Okol-Glava, Gniljane, Sultan (Nevski) Popovo, and Settlement mound Burgas); Late Chalcolithic 6500-6000 BP (Dolnoslav, Varna); Bronze 6000-5000 BP (Urdoviza – Kiten, Baley); Late antiquity (Kapitan Andreevo, Dyadovo village – Nova Zagora, Charda village, Yambol district, Trakia motorway, Yambol district, and Academic, Plovdiv) (S1 Table and S1 Fig).

The *Canis familiaris* remains were determined and assigned to the two primitive prehistoric morphotypes (and probable crossbreeds between them), representing the two main Mesolithic\Neolithic-Chalcolithic canine races of the Balkans that can be called conditionally *Canis familiaris* “*intermedius*” and *C. f*. “*palustris*”. The mesolithic form “*vlasac*”, described from Serbia [30] is not different from *C. f. intermedius* and represents a slightly younger form of the latter, with skull/tooth dimensions that are in general within the lower values of the individual variations of *C. f. vlasac*, while “*C. f. palustris*” is the more domesticated small form [31].

Ancient DNA isolation and PCR amplification and sequencing ware given in the Supporting Information S1.

### Phylogenetic reconstruction

The obtained sequences were manually edited and aligned by MEGA software version 7.0^32^, using the dog mtDNA sequence NC_002008 [33] and EU789787 [8] as a reference. Sequences were analyzed by polymorphic SNPs, and haplogroups were determined according to [9] and [34] as well as MitoToolPy program [35], (http://www.mitotool.org/mp.html) with reference sequence EU789787 [8]. The phylogenetic analysis was based on the archaeological dog samples used in this study as well as on all available in GenBank ancient DNA dog sequences [5,12–14,16–18,21,27–29,36]. Ancient and recent wolves [5,21–24,37], and the recent Bulgarian native dog [38] were characterized using network analysis – NETWORK 4.5.1.6 (Fluxus Technology Ltd.) (available at http://fluxusengineering.com).

In order to graphically display (and summarize) the mitochondrial relationships among the analyzed ancient dog populations and all ancient samples available in GenBank, we performed a principal component analysis (PCA) – a method that considers each haplogroup as a discrete variable and allows a summary of the initial dataset into principal components (PCs). Principal component analyses (PCA) were performed using Excel software implemented by XLSTAT, as described elsewhere [39]. The PCA were carried out considering all available ancient dog sample worldwide.

The obtained sequences included in this study were deposited in the National Center for Biotechnology Information (NCBI) GenBank database under accession numbers NCBI: MH937186–MH937206.

### Data availability statement

Consensus sequences from this study are available from the GenBank database (accession numbers MH937186–MH937206).

### Author Contributions

M.M., G.R., and P.H. conceived of and designed the experiments; G.R., P.H., and N.S. wrote the manuscript text; I.Y. and M.P. performed the experiments; G.R., P.H., M.M., and N.S. analyzed the data.

### Competing interests

The authors declare no competing interests.

